# Highly multiplexed fluorescent imaging via fluidic-free, rapid sequential plasmonic heating activated signal exchange of DNA probes

**DOI:** 10.64898/2026.02.04.703891

**Authors:** Ethan Xu, Yanju Chen, Qing Wang, Aadhya A. Harugeri, Md Yeasin Pabel, Rachel S.F. Moor, Qianqian Song, Elias Sayour, Ashley Parham Ghiaseddin, Wei David Wei, Fan Hong

## Abstract

Fluorescence microscopic methods are critical for spatial profiling of multiple biological targets in cells and tissues to study cell and tissue functions, but their multiplexity was limited to 3∼5 targets under a conventional setup using different fluorescent channels because of spectra overlap. Here, we introduce a simple, rapid, wash-free multiplexed fluorescence imaging method in cells and tissues, termed DNA based plasmonic heating activated signal exchange reaction (PHASER). PHASER uses infrared light-induced plasmonic heating of gold bipyramid nanoparticles to sequentially activate thermodynamically calibrated DNA thermal probes in situ for rapid and multiplexed fluorescent imaging of biological targets. We showed that the signal exchange per round between biological targets in PHASER can be completed within 30 seconds, and that 5 irradiation pulses of photothermal heating can activate DNA thermal probes with 5 different signal temperatures in cells and tissues. To demonstrate its practical use, we applied PHASER to profile the subcellular spatial organization of different organelles in cultured cells and validated its performance in FFPE tissue. We further used PHASER to resolve up to 15 different protein spatial expression profiles in mouse brain tissue with dimensions of millimeters for single-cell proteomic analysis. PHASER is expected to have broad biotechnical applications with multiplexed fluorescence imaging for a wide variety of biological targets across diverse samples.

## INTRODUCTION

Fluorescent imaging has been widely used to visualize biomolecules in cells and tissues to study their biological function. Biological targets are usually bound to a fluorescent-labeled binding entity to reveal the target’s quantity and spatial location within the sample in situ. For example, the immunofluorescence method uses fluorophore-labeled antibodies to image proteins in cells and tissues with subcellular resolution, enabling the study of cellular function and disease mechanisms with phenotype information.^1–3^ To obtain higher-dimensional biomolecular information to get deep molecular insights, multiplexed fluorescence imaging of different targets is required.

Conventional fluorescent microscopy methods usually use spectrally separated fluorophores, which are limited to 3∼5-plex due to spectral overlap. While hyperspectral properties of dyes can enable higher-plex imaging^4^, this requires complex detectors and algorithms to resolve signals. Sequential imaging methods (for example, cycIF^5, 6^, 4i^7^, and COMET^8^) overcome this limitation by switching signals between targets through iterative labeling of antibodies, imaging, and bleaching. However, the overall imaging workflow becomes very long due to the antibody staining and stripping, each round takes hours to complete. Later, to avoid the lengthy iterative process of primary antibody staining, DNA-based barcoding and exchange methods (for example, CODEX^9, 10^, DEI^11, 12^, and SABER^13, 14^) have been developed in which all targets are stained in a single step with DNA-barcoded antibodies, followed by iterative binding and washing of fluorescently labeled DNA imagers. As the DNA imagers used are usually 15-20-nt long and much smaller than the bulky antibodies, the signal exchange step with DNA is much faster and easier. These approaches have greatly expanded multiplexed protein imaging.^15^ However, because the fluorescence signal is carried by newly introduced chemical labels, each imaging round still requires reagent delivery, incubation, washing or signal erasure through fluidic exchange (**Supplemental Table S1**). The resulting fluidic workflows are slow, instrumentally complex and increasingly difficult to implement in large or thick tissue specimens, where reagent transport is diffusion limited. A new chemical mechanism for signal-switching that operates directly within the specimen, without repeated fluidic exchange-based washing steps, would make multiplexed protein imaging faster and more accessible.

In conventional fluorescence imaging, light functions primarily as a readout, excitation wavelengths are matched to spectrally distinct fluorophores to preserve signal orthogonality. This wavelength–color pairing inherently limits multiplexity. We asked whether light could instead act as a programmable chemical input, enabling multiple targets labeled with the same fluorophore to be activated or erased according to probe state rather than dye color. To achieve this, light would need to serve as a chemical input that controls probes carrying the same fluorophore in different molecular states, rather than only as an optical input that excites fluorescence. In this design, a single wavelength of light would not encode target identity through dye color but would instead be converted into programmable reaction conditions that activate or erase fluorescence from molecular probes in situ. Other than the increase of multiplexed imaging capacity, the photothermal strategy enables rapid and convenient signal activation on a standard fluorescence microscope with minimal and low-cost set-up.

Here we introduce plasmonic heating-activated signal exchange reaction, or PHASER, a light-controlled chemical strategy for multiplexed protein imaging. PHASER uses gold bipyramidal nanoparticles dispersed within the sample to convert infrared light into calibrated photothermal pulses. These pulses create defined temperature inputs that control the hybridization state of thermodynamically programmed DNA thermal probes bound in situ to DNA-barcoded antibodies. At lower thermal thresholds, the quencher strands dissociate, activating fluorescence; at higher thresholds, the reporter strands dissociate, erasing the signal. Thus, PHASER converts optical irradiation into programmable DNA strand-exchange reactions directly within fixed cells and tissues, enabling rapid, light-controlled signal switching without repeated fluidic exchange. Compared with Thermal-plex imaging,^16^ which relies on electronic heating with specialized heating station and sample holding chamber, PHASER provides a more readily controlled and implementable light-based approach for thermal activation. It also has potential for more localized heating activation in specific region or even heating in volumetric tissues.

To establish PHASER for multiplexed protein imaging, several chemical and bioanalytical challenges must be addressed. DNA-barcoded antibodies must retain antigen-binding activity during repeated photothermal cycles. Gold bipyramids must remain colloidally stable in immunostaining and imaging buffers, produce reproducible temperature pulses, and avoid interfering with fluorescence readout. DNA thermal probes must preserve well-separated melting thresholds and efficient signal activation and erasure across diverse targets and specimen types. We therefore systematically validated nanoparticle stabilization, infrared pulse calibration and thermally programmed signal switching in cultured cells, human FFPE tissue and fresh-frozen mouse brain sections. Using PHASER, antibody-bound DNA thermal probes were activated within 30 s, and five infrared pulse conditions selectively addressed thermal channels at 39, 48, 57, 65, and 72 °C. We demonstrate multiplexed imaging of subcellular organelle markers in cultured cells, selective DNA thermal probe activation in clinically relevant FFPE and fresh-frozen tissues, and high-plex spatial protein profiling across millimeter-scale mouse brain sections. The high-plex spatial proteomics resolved by PHASER enables single-cell proteomic analysis for cell typing and mapping to study tissue organization. Together, these results establish PHASER as a rapid, light-controlled, and fluidics-free platform for highly multiplexed protein imaging and, more broadly, as an optical approach for programming DNA reaction states in biological specimens.

## Results

### Principle of PHASER

The principle of PHASER is to use AuBPs to convert IR irradiation into heat via plasmonic heating and thereby activate DNA thermal probes bound to targets in situ. Plasmonic heating arises when nanomaterials convert light energy into heat through collective electron oscillations^17^. The absorption of light by plasmonic nanostructures and the resulting temperature increase are exquisitely sensitive to the structure’s shape and composition and to the illumination wavelength^18^. Gold bipyramidal nanoparticles (AuBPs) exhibit strong surface plasmon resonance and efficiently generate heat when irradiated with 800–900 nm light^19^ and has been used in photothermal therapy^20, 21^ and bioanalysis^22^, owing to their high aspect ratio (longitudinal length to side width). The response wavelength range has minimal interference with the fluorescent channels typically used in immunofluorescence imaging, making IR irradiation a non-interfering resource for the fluorescent imaging.

To image a target of interest in situ, cells and tissue samples undergo a standard preparation process that includes fixation, permeabilization, and staining. As shown in **Figure 1a**, a primary binding entity (e.g., in situ hybridization probes, antibodies, aptamers) labeled with a unique DNA barcode is used for sample staining. This is followed by the binding of complementary DNA thermal probes and subsequent washing steps. Each DNA thermal probe is composed of a quencher strand labeled and a reporter strand designed to hybridize with the DNA barcode in a temperature-dependent manner. The quencher binding domain’s melting temperature T_mq_ is designed to be 6∼8 °C lower than the imager binding domain T_mb_. Therefore, at the signal temperature T_s_, which is between the T_mq_ and T_mb_, the quencher is melted off for fluorescence activation, whereas the reporter strand remains bound until exposure to a higher temperature, at which point it is also released and the signal is erased. This differential melting design enables programmable on/off fluorescence switching using brief heating spikes at defined temperature channels, and 5 different temperature channels have been developed previously.^16^ After dispersing gold bipyramids (AuBPs) in the sample, a precisely calibrated infrared (IR) pulse is applied to induce a rapid heating spike via plasmonic resonance effects. This heating selectively activates the fluorescent signal of the bound DNA thermal probes by melting off the quencher strand. Imaging is then performed. Subsequently, longer or stronger IR pulses are applied to melt the imager (reporter) strand, thereby removing the fluorescent signal.

**Figure 1.**
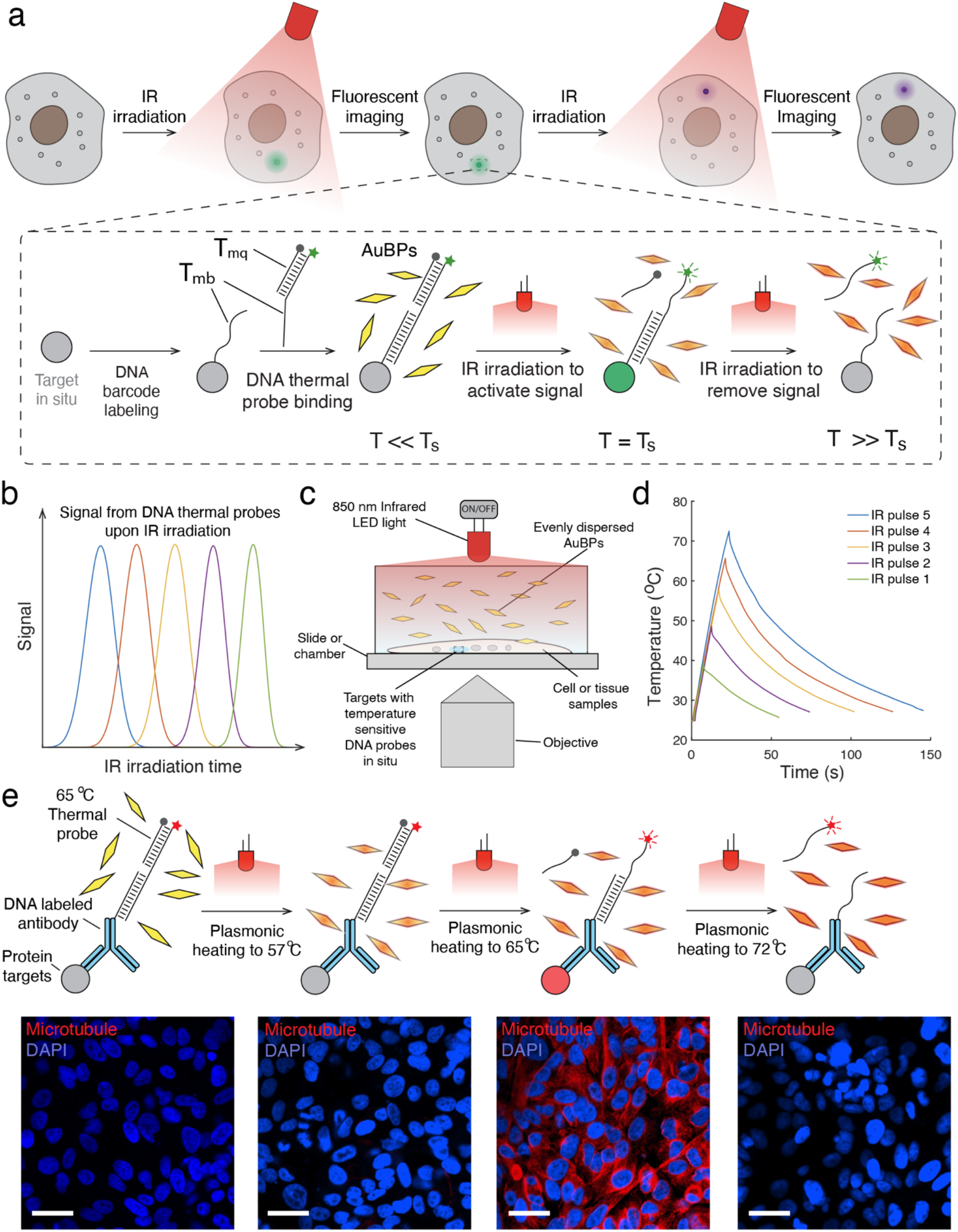
Principle of PHASER. **(a)** The schematics of PHASER. The target of interest will be labeled with a DNA barcode. The DNA thermal probe is consisted of imager and quencher, and the quencher binding domain’s melting temperature Tmq is designed to be 6∼8 °C lower than the imager binding domain Tmb. At the thermal probe’s designed signal temperature of Ts, the imager remains binding to the barcode in situ and quencher is melted off for PHASER imaging. After the addition of gold bipyramid nanoparticles, the infrared LED light is applied to the sample for plasmonic heating to generate heating spike and activate the DNA thermal probe’s signal. After the fluorescent imaging of the sample, another longer pulse of infrared light irradiation is applied to melt off the imager and remove the signal from the targets. Multiple rounds of plasmonic heating can be applied for multiplexed imaging. **(b)** The relationship between the fluorescent signal of DNA thermal probes with different signal temperatures and the IR irradiation time. The fluorescent signal of thermal probes can be activated orthogonally with calibrated IR irradiation time. **(c)** The set-up of PHASER protein imaging. **(d)** The 5-temperature spike generated from calibrated infrared light irradiation induced plasmonic heating for the activation of DNA thermal probes with 5 different signal temperatures. **(e)** The example of alpha tubulin imaging in cultured U2OS cells with photothermal stimulation of DNA thermal probes at 65 °C. All the scale bars are 40 µm.

The DNA thermal probes are designed with five distinct and well-separated signal temperatures, enabling unambiguous multiplexing across five different temperature spikes. Five sequentially calibrated IR irradiation pulses are used to match these thermal channels through controlled plasmonic heating (**Figure 1b**). Multiplexed imaging is achieved in a highly streamlined manner. All DNA barcoded binding entities are applied in a single staining step, followed by simultaneous binding of all DNA thermal probes. Sequential IR pulses then activate and subsequently erase the fluorescent signals of the probes in situ, allowing sequential imaging rounds. The resulting multi-round fluorescence images are computationally aligned and overlaid to generate multi-channel images for each target.

### Validation of PHASER imaging

We first used PHASER to visualize protein targets in cultured cell samples for the concept validation. To implement the plasmonic heating for multiplexed protein imaging, a simple, low-cost IR LED light was used to irradiate AuBPs dispersed throughout the sample, generating even heating regardless of sample dimensions when the IR light can cover the sample (**Supplemental Figure S1**). IR light from an LED was positioned above the sample and induces plasmonic heating (**Figure 1b**). A thermocouple was inserted into the sample chamber to monitor temperature for calibration. We first validated absorbance of AuBPs and tested the influence of different concentrations (1, 5, and 10 OD) of AuBPs on the plasmonic heating. We found a concentration of 1 OD in the sample buffer already achieved rapid heating for the highest thermal channel with 50W of LED irradiation with about 23 seconds, and 5 and 10 OD didn’t show significant increase of heating speed as shown in **Supplemental Figure S2**). We then calibrated IR pulse duration against the resulting heating spikes with 1 OD AuBPs, and the activation temperatures of the five developed thermal probes are reached in 7.5 s, 12.6 s, 16.8 s, 20.5 s, and 22.6 s (**Figure 1d**). We further tested the compatibility of AuBPs with in situ fluorescent imaging conditions. The AuBPs was found to aggregate in 1x PBS buffer, and their absorbance in 850 nm disappeared (**Supplemental Figure S2**) due to the high salt concentration. BSA was then used to coat the synthesized AuBPs^23^ to enhance compatibility with the in situ imaging environment and successfully prevented the aggregation in 1x PBS buffer (**Supplemental Figure S2**).

We then moved to test the performance of protein imaging with PHASER, an alpha-tubulin antibody was conjugated with a DNA barcode to through click chemistry^24^ (**Supplemental Figure S3**). Briefly, a DBCO-NHS crosslinker was first applied to link to the amine residue on the antibody and then conjugated to an azide-labeled DNA barcode. If the amine residue conjugation failed, DBCO-Maleimide crosslinker were applied to cysteine residue. Cultured U2OS cells were used for the initial validation. After penetration and fixation of cultured U2OS cells, the DNA-conjugated alpha-tubulin antibody was used to stain the cell samples. Then, a DNA thermal probe labeled with an Atto647–BHQ3 pair and a signal temperature of 65 °C was applied to bind the antibody (See probe information **Supplemental Table S2** and antibody information in Supplemental **Table S3**). The overall workflow (**Supplemental Figure S4)** dramatically simplifies multiplexed in situ protein imaging compared to traditional methods. As shown in **Figure 1d**, BSA-treated AuBPs were suspended in the imaging buffer and added to the sample for PHASER imaging. No alpha tubulin signal was observed at room temperature, indicating minimal influence of AuBPs on imaging and efficient quenching. Application of a calibrated infrared pulse 3 to irradiate the AuBPs for plasmonic heating yielded no detectable signal. After infrared pulse 4, the alpha-tubulin signal appeared clearly. When a longer infrared pulse 5 was applied, the alpha-tubulin signal was removed.

### Performance evaluation of PHASER

We then evaluated the speed of signal activation and removal in PHASER protein imaging. As shown in **Figure 2a** and **2b** for the fluorescent images and single cell protein signal level, 30 seconds of plasmonic heating with infrared pulse 4 fully activated the signal with an S/N of 3.34 ± 0.77, and an additional 60 seconds of heating did not further significantly improve it. Signal removal of DNA thermal probes via plasmonic heating was also rapid and was completed within 30 seconds (**Figure 2c** and **2d**). The S/N was reduced from 4.06 ± 0.83 to almost background level. After the dissociation of the DNA imagers and quenchers, these strands diffused into the imaging buffer and remain very low concentration (fM to pM range). Therefore, they are unlikely to rebind to the original antibody barcode due to extremely slow kinetics.^16^ We noticed the background signal before plasmonic heating (S/N = 1.12 ± 0.05) is slightly higher than the background after plasmonic heating to 72 °C (S/N = 1.01 ± 0.02), which may be due to the incomplete quenching of the fluorophores. The previously reported quenching efficiency is about 95 to 98%.^25^ Other fluorescent background may come from non-specific binding of antibody or fluorescent imagers, and heating of samples can help to reduce fluorescent background by facilitating the dissociation of unspecific binding. Overall, these results demonstrated the unambiguous and rapid signal switching of using PHASER for protein imaging, and off-thermal channel signal is almost background level.

**Figure 2.**
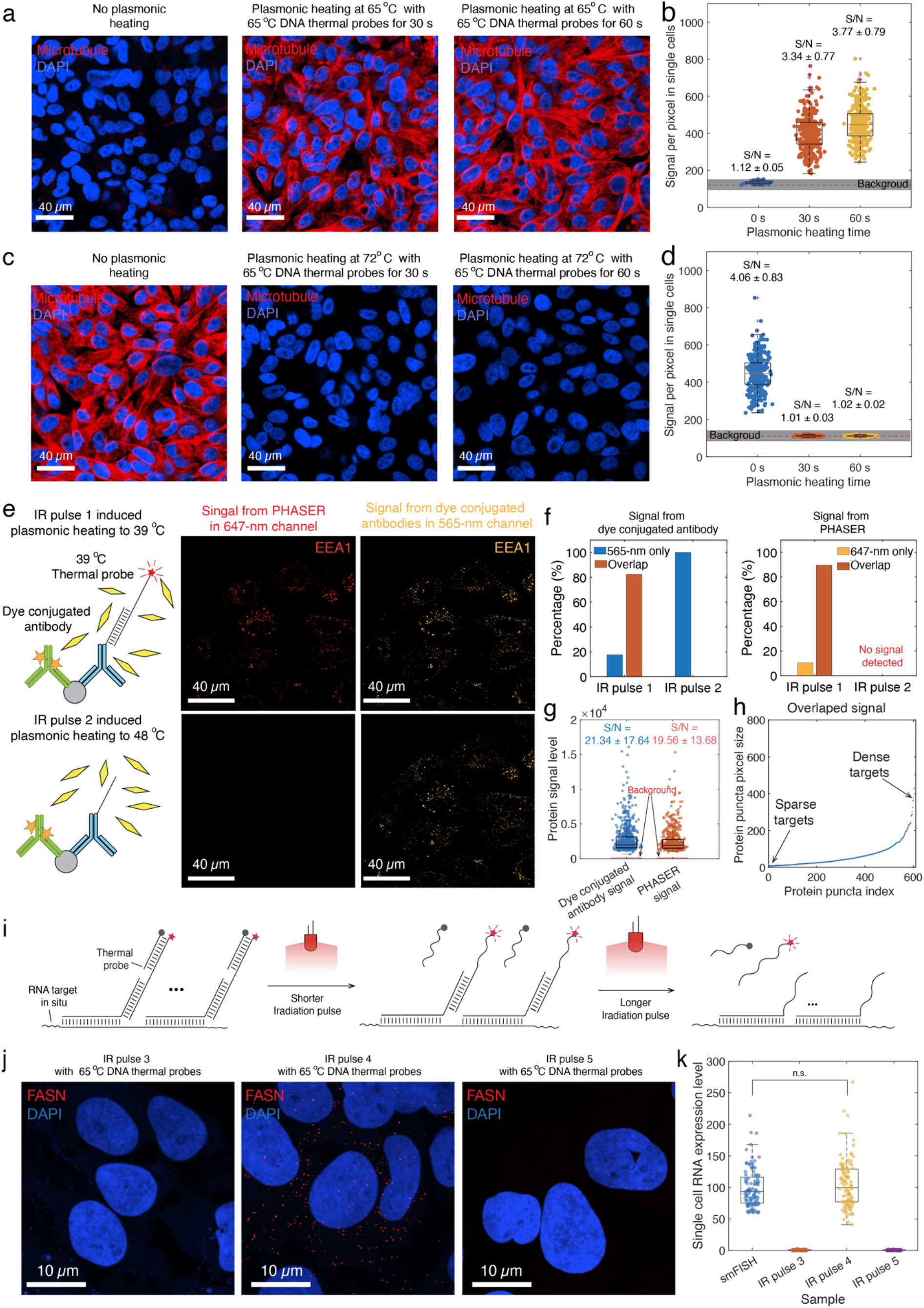
Evaluation of PHASER performance and its broad target applicability. (**a**) Representative microtubule images obtained with 65 °C DNA thermal probes before heating and after plasmonic heating at 57 °C for 30 s or 65 °C for 60 s. (b) Corresponding single-cell fluorescence intensities and signal-to-noise ratios (S/N). The cell numbers used are 244, 355, and 318, respectively. (**c)** Microtubule images before and after probe removal by plasmonic heating at 72 °C for 30 or 60 s. (**d)** Corresponding fluorescence intensities and S/N values. The cell numbers used are 327, 500, and 500, respectively. (e) Specificity analysis of EEA1 detection using a DNA-conjugated antibody imaged by PHASER in the 647-nm channel and a fluorophore-conjugated antibody targeting a different epitope in the 565-nm channel. PHASER signals appeared after heating to 39 °C and were removed after heating to 48 °C, whereas the fluorophore-conjugated antibody signal was retained. (**f)** Fractions of puncta detected only at 565 nm, in both channels and only at 647 nm following each IR pulse. Of the PHASER puncta, 89.4% overlapped with fluorophore-antibody puncta, and 83.4% of fluorophore-antibody puncta overlapped with PHASER puncta. (**g**) The comparison activated signal of PHASER with fluorophore conjugated antibody for the same protein targets in situ. (**h)** Size distribution of EEA1 puncta detected in both channels. (**i)** Schematic of irradiation-duration-dependent control of thermal-probe hybridization and dissociation. **(j)** Representative FASN RNA images obtained with 65 °C DNA thermal probes after sequential plasmonic heating at 57, 65 and 72 °C. (**k)** Single-cell FASN RNA levels measured by smFISH and after the indicated IR pulses. There is no significant difference between the resolved single cell RNA expression from smFISH and PHASER with IR pulse 4 (p value = 0.22, two tail t test). Scale bars, as indicated. N>3 replicates were conducted for all the experiments.

Unlike bulk protein analysis in vitro, the fluorescence intensity generated by PHASER for in situ protein imaging depends strongly on the local abundance of target proteins and the binding affinity of the corresponding antibody. To further evaluate the specificity and sensitivity of PHASER, we compared PHASER signals detected in the 647-nm channel with signals from a fluorophore-conjugated antibody detected in the 565-nm channel, using EEA1 as the target. EEA1 is a well-established marker of early endosomes and was selected because of its characteristic punctate, approximately spherical distribution. The broad range of endosome sizes also reflects variations in local target abundance, making EEA1 a suitable marker for assessing the specificity and sensitivity of PHASER.

The DNA-conjugated antibody for the DNA thermal probe in the 647-nm channel and the fluorophore-conjugated antibody for the 565-nm channel will target different epitopes of EEA1 (**Supplemental Table S4**). As shown in **Figure 2e**, the PHASER signal appeared in the 647-nm channel following plasmonic heating induced by the IR pulse 1 and disappeared after heating induced by the IR pulse 2. In contrast, the signal from the fluorophore-conjugated antibody in the 565-nm channel remained consistently detectable. Among the puncta detected in the 647-nm channel, 89.4% overlapped with puncta in the 565-nm channel; conversely, 83.4% of puncta in the 565-nm channel overlapped with those in the 647-nm channel (**Figure 2f** and **Supplemental Figure S5**). This high degree of colocalization supports the specificity of PHASER regardless of nonspecific binding from the antibody. The non-colocalized puncta are much smaller (**Figure S5**), likely due to nonspecific background from both antibodies. Comparison of the protein signal and background levels from PHASER and fluorophore-conjugated antibody labeling of the same protein targets in situ showed comparable signal and background distributions between the two methods, resulting in similar levels of S/N ratios (**Figure 2g**). To further evaluate sensitivity, we further analyzed the areas of the colocalized puncta (**Figure 2h**). Their sizes ranged from 6 to 640 pixels, demonstrating that PHASER detects EEA1-positive structures across a broad range of local target abundances with sensitivity comparable to that of conventional fluorophore-conjugated antibody imaging. Additional specificity data comparison is shown in **Figure S6** with alpha microtubule imaging (**Supplemental Table S4** for antibody information).

To demonstrate PHASER’s broader applicability beyond proteins, we further validated its performance for RNA imaging (**Figure 2i**). We targeted FASN mRNA in cultured U2OS cells using 65 °C thermal probes that respond to IR pulse 4-induced plasmonic heating-mediated activation. The in-situ hybridization DNA probes were designed using Oligominer^26, 27^ with melting temperatures above 75 °C to avoid dissociation during imaging (see sequence information in **Supplemental Table S5**). The thermal probe showed minimal FASN RNA signal under IR pulse 3 and IR pulse 5, whereas IR pulse 4-induced plasmonic heating recovered robust FASN RNA signals (**Figure 2j**). Quantification of single-cell RNA expression levels confirmed selective detection of the target RNA at the intended infrared irradiation pulse, with low background expression estimates under off-target pulse conditions. The resolved single-cell RNA expression levels were also consistent with smFISH-resolved RNA measurements, demonstrating the accuracy of the method (**Figure 2k**). These results demonstrate that plasmonic heating can selectively activate RNA probes in situ and enable temperature-controlled RNA expression readout at the single-cell level.

### Validation of 5 IR irradiations with PHASER imaging

We next validated signal activation for five DNA thermal probes with activation temperatures of 39, 48, 57, 65, and 72 °C using the five IR pulses by imaging alpha tubulin in fixed U2OS cells. We first validated the performance of DNA thermal probes for alpha tubulin protein imaging by electronically heating each to all the 5 temperature channels. All 5 DNA thermal probes successfully activated at their assigned thermal channels and remained non-detectable in other channels (**Supplemental Figure S7**), confirming the robustness of DNA thermal probes for protein imaging. The corresponding IR pulses for each thermal channel were then used to activate the DNA thermal probes with AuBPs dispersed in situ. As shown in Figure 3a, for each DNA thermal probe, application of its matched IR pulse specifically activated the signal, whereas shorter or longer pulses did not generate any signal (**Figure 3a**). We also analyzed single-cell alpha tubulin signals based on the fluorescent images, and cells receiving matched IR pulses were found to have significantly higher signals than those receiving shorter or longer pulses (**Figure 3b**). Together, the fluorescence images and single-cell protein signal analysis (see Methods and **Supplemental Figure S8**) demonstrated the robustness of using five IR pulses to specifically activate the fluorescence signal of the corresponding five thermal probes for protein imaging. **Multiplexed imaging to resolve subcellular distribution of organelles with PHASER in a single fluorophore channel.** The robustness of signal activation using plasmonic heating across five thermal probes with distinct activation temperatures enables rapid and fluidic-exchange-free multiplexed fluorescence protein imaging. To demonstrate this, we selected five protein targets, including alpha-tubulin, clathrin, cis-Golgi matrix protein 1 (GM130), a mitochondrial marker, and early endosomal antigen 1 (EEA1), to visualize five organelles, including microtubules, vesicles, Golgi, mitochondria, and early endosomes, respectively. These organelles have diverse functions, including cytoskeletal support, intracellular transport, protein and lipid processing, energy generation, and cargo sorting. Antibodies targeting these proteins were conjugated to five different DNA barcodes (**Supplemental Table S2 and S3** and **Supplemental Figure S9**), allowing binding to five DNA thermal probes with activation temperatures of 39, 48, 57, 65, and 72 °C. All five DNA thermal probes were encoded with an Atto647–BHQ-3 pair, enabling the use of a single fluorophore channel to image five protein targets.

**Figure 3.**
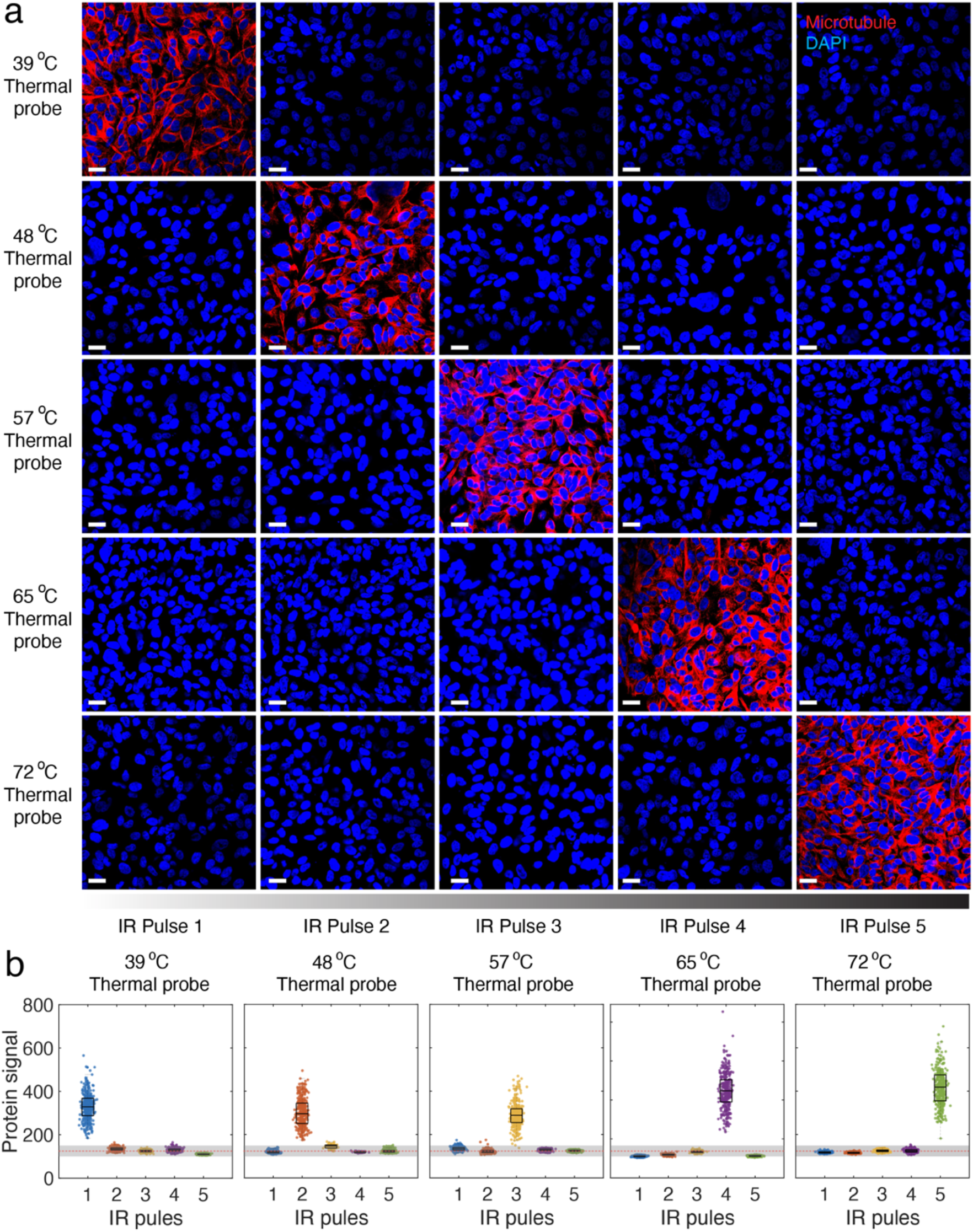
Validation of PHASER using five DNA thermal probes and five infrared (IR) pulse conditions in cultured cells. **(a)** Representative immunofluorescence images of α-tubulin acquired after plasmonic heating under five IR pulse conditions using five DNA thermal probes with calibrated activation (signal) temperatures of 39, 48, 57, 65, and 72 °C. Scale bars, 20 µm. **(b)** Single-cell α-tubulin signal intensity per pixel for each thermal probe (39, 48, 57, 65, and 72 °C, respectively) across the five IR pulse conditions. The counted cell numbers are ranging from 224 to 500 cells. Boxplots summarize the distribution (median and interquartile range; whiskers, 1.5× interquartile range). The shaded gray region and red dashed line indicate the background signal and its standard deviations. N >3 replicates were conducted for all the experiments.

After DNA barcode conjugation and validation, we first tested the DNA thermal probes bound to each antibody individually. No significant crosstalk between thermal channels was observed (**Supplemental Figure S10**). We then mixed all five DNA-barcoded antibodies and stained fixed U2OS cells in a single step (**Figure 4a**). A mixture of five DNA thermal probes was applied, and excess probes were washed away. The sample was then imaged with plasmonic heating after addition of AuBPs suspended in imaging buffer. Five rounds of IR pulse–induced plasmonic heating and imaging were applied to sequentially activate fluorescence for the five proteins. As shown in **Figure 4b**, all five protein signals are clearly resolved at subcellular resolution. **Figure 4c** shows their overlap with DAPI, clearly depicting organelle distributions.

**Figure 4.**
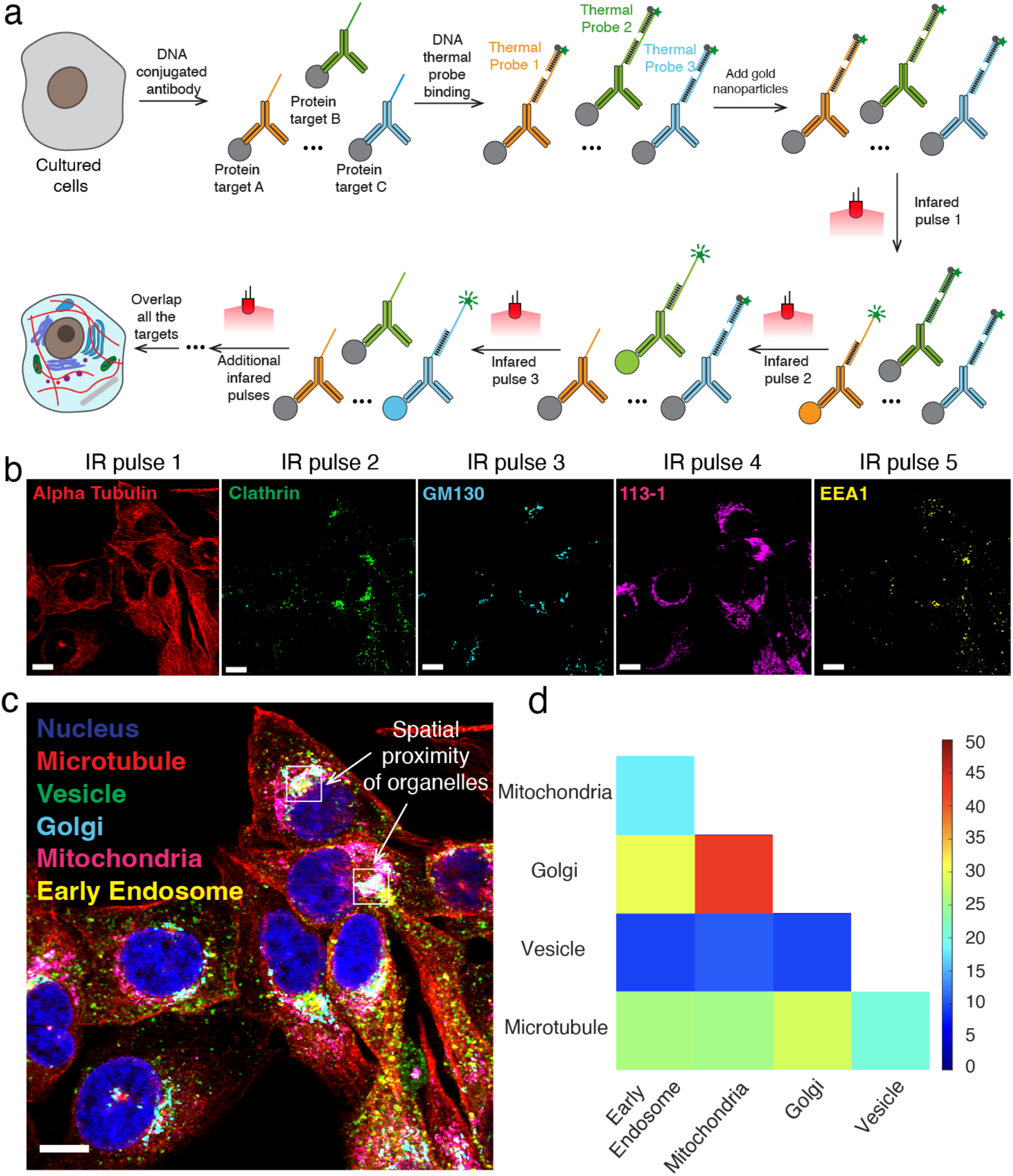
PHASER enabled multiplexed imaging in cultured cells with a single fluorophore channel. **(a)** Schematics of multiplexed fluorescent imaging with PHASER. **(b)** The individual images of each protein marker, including alpha tubulin, clathrin, GM130, 113-1, and EEA1, in fixed U2OS cells. **(c)** The overlapped images of different organelles were resolved by the protein markers. Boxed regions indicate the physical proximity of the vesicle with Golgi. **(d)** Quantitative analysis of organelle proximity based on median distance between the centroids of resolved organelle. Color bar indicates the median pixel distance. All the scale bars are 10 µm. N >3 biological replicates were conducted for all the experiments.

These multiplexed data, enabled by PHASER’s rapid imaging of multiple organelles, provide insight into how they communicate to maintain cellular function.^7, 28, 29^ We observed substantial signal overlap between transport organelles, vesicles (clathrin), endosomes (EEA1), and the Golgi (GM130). These spatial signal proximity support the role of the Golgi in sorting and packaging modified proteins and lipids into transport vesicles for delivery to other cellular sites. The overall spatial distribution relationship between all the 5 organelles is shown as the median distance between the centroids of resolved cellular organelles in **Figure 4d**. With more fluorophore channels integrated and higher multiplexity achieved, PHASER could image additional organelles and help unravel cellular molecular complexity in studies of cell trafficking and drug response.

**Validation of PHASER in** human FFPE and fresh-frozen **mouse brain tissues.** Immunofluorescence imaging of protein markers in tissue sections is widely used in neuroscience, cancer research and clinical pathology.^30–32^ To evaluate whether PHASER is compatible with commonly used tissue formats, we tested probe activation in both human formalin-fixed paraffin-embedded (FFPE) tissue and fresh-frozen mouse brain sections. We first examined PHASER performance in human tonsil FFPE tissue using a DNA conjugated CD45 antibody (**Supplemental Table S4**) bound with 57 °C DNA thermal probe. In the absence of plasmonic heating, CD45 fluorescence was efficiently quenched, resulting in minimal background signal. A subthreshold IR pulse that heated the sample to 48 °C also produced little CD45 signal. In contrast, the matched IR pulse that generated a 57 °C heating spike robustly activated CD45 fluorescence. Subsequent heating to 65 °C removed the signal, consistent with thermal displacement of the reporter strand from the DNA barcode (**Figure 5a**). Line-scan intensity profiles across the same regions confirmed selective activation at 57 °C and signal removal after the higher-temperature pulse (**Figure 5b**). These results demonstrate that PHASER can operate in FFPE tissue, a sample format broadly used in pathology and clinical archives.

**Figure 5.**
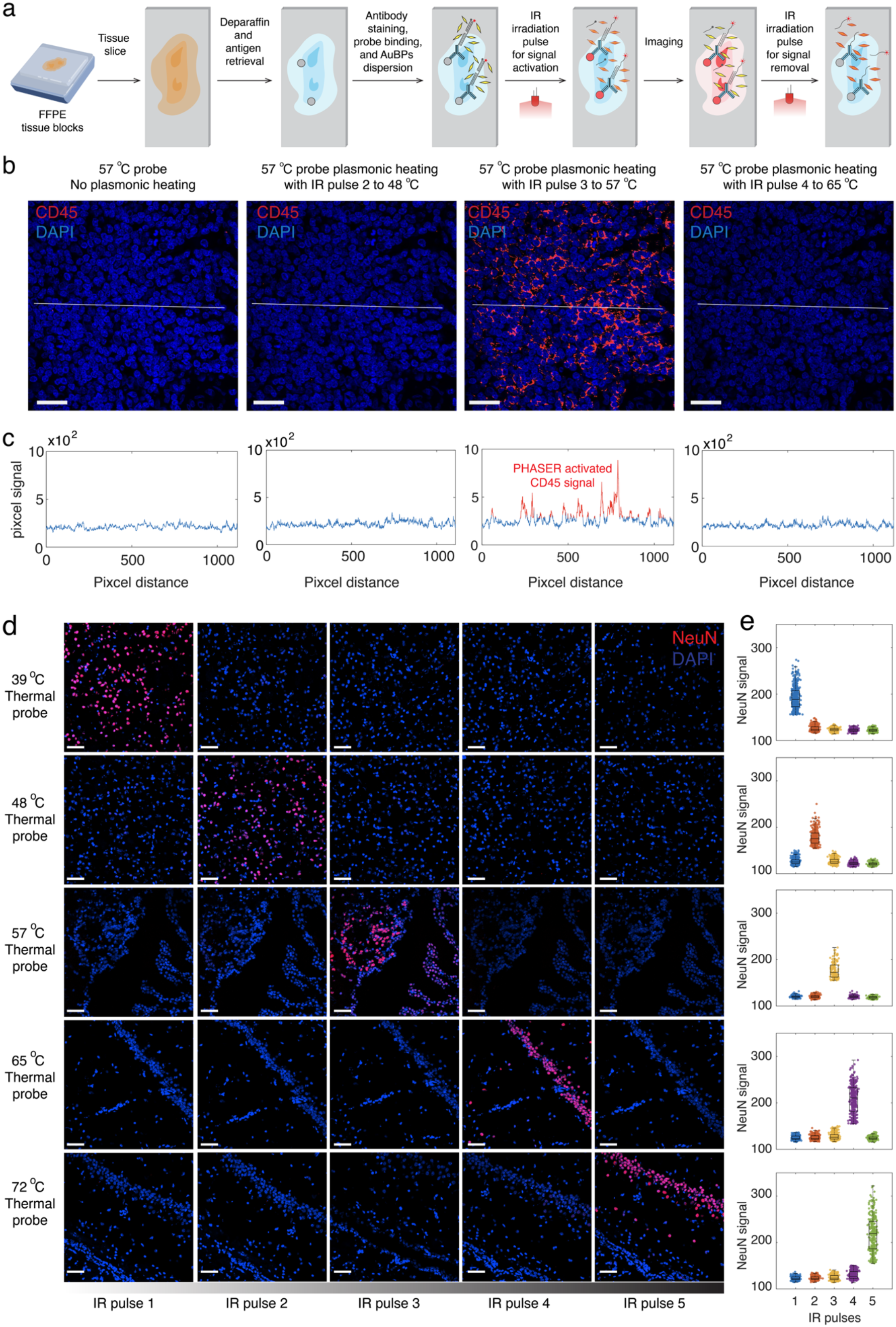
Validation of PHASER in FFPE human and fresh-frozen mouse brain tissue sections. **(a)** The workflow of PHASER imaging for FFPE tissues. (**b)** Representative fluorescence images of human tonsil FFPE tissue stained with a CD45 and 57 °C DNA thermal probes. Images were acquired without plasmonic heating or after AuBP-mediated plasmonic heating with IR pulse 2, pulse 3 or pulse 4, corresponding to heating spikes of 48, 57 and 65 °C, respectively. Minimal CD45 signal was detected without heating or after the subthreshold 48 °C pulse, whereas robust CD45 signal was activated by the matched IR pulse 3 to 57 °C. The signal was removed after longer IR pulse 4 to higher-temperature 65 °C pulse. Scale bars, 50 µm. (**c)** Line-scan fluorescence intensity profiles measured along the lines indicated in **b**. The profiles show low background signal before activation and after the 48 °C pulse, strong CD45 signal in red after the matched 57 °C pulse and signal removal after heating to 65 °C. (**d)** Representative fluorescence images of fresh-frozen mouse brain tissue stained with NeuN DNA thermal probes calibrated to five thermal channels: 39, 48, 57, 65 and 72 °C. Rows indicate the activation temperature of each NeuN thermal probe and columns indicate the applied IR pulse condition. For each probe, NeuN fluorescence was selectively activated by the matched IR pulse, with minimal signal detected under non-matched pulse conditions. Scale bars, 50 µm. (**e)** Single-cell quantification of NeuN fluorescence intensity from **d** across the five IR pulse conditions for each thermal probe. N >3 biological replicates were conducted for all the experiments. Points represent individual cells; box plots show the median and interquartile range, with whiskers extending to 1.5× the interquartile range. The cell populations for all conditions are n > 300 cells. The matched pulse condition produced the highest NeuN signal for each thermal channel, confirming selective probe activation in fresh-frozen tissue.

We next evaluated PHASER in fresh-frozen mouse brain sections, a structurally complex tissue with dense cellular organization. Mouse brain slices with a thickness of ∼10 µm were prepared on a cryostat. NeuN ^33, 34^, a neuron-specific nuclear protein that marks mature, post-mitotic neurons in the vertebrate nervous system, was chosen for validation of signal activation and removal with all 5 IR irradiation pulses. An antibody targeting NeuN was conjugated to DNA barcodes and validated in situ before PHASER imaging (**Supplemental Figure S11**). Five IR pulses were applied to samples labeled with five thermal probes, and fluorescence imaging was used to evaluate the NeuN signal in brain tissue. Fluorescence images used to evaluate the performance of the five IR pulses were acquired from different regions of the brain slices. As shown in **Figure 5c**, the NeuN signal appeared only when matched IR pulses were applied, indicating PHASER’s robustness in complex tissue samples. Single-cell NeuN signal was quantified by average per-cell pixel intensity (**Figure 5d**), and the NeuN signal in mature neuronal cells was significantly higher than in cells exposed to non-matched pulses. Together, these results establish that PHASER enables thermally programmed, target-specific signal activation and removal in both human FFPE tissue and fresh-frozen mouse brain tissue. The successful validation across these distinct tissue preparations supports the robustness of PHASER for spatial protein imaging in biologically and clinically relevant samples.

### PHASER enables scalable and 15-plex multiplexed protein imaging in mouse brain tissues

Spatial analysis of brain tissue requires the simultaneous measurement of multiple protein markers that define cell identity, tissue architecture and functional state.^30, 35^ To test whether PHASER could support high-plex protein imaging across a millimeter-scale tissue region, we applied the method to coronal mouse brain sections containing hippocampal and adjacent brain structures. We selected 15 protein markers that report major neural and glial cell types, neuronal projections, synaptic structures and cytoskeletal organization: tubulin beta-3 (TUBB3), neuropeptide Y (NPY), glial fibrillary acidic protein (GFAP)^36, 37^, vesicular glutamate transporter 1 (VGLUT1), 2’,3’-cyclic-nucleotide 3’-phosphodiesterase (CNPase) and Synaptophysin (SYP), Myelin basic protein (MBP), Ribosomal protein S6 (RPS6) and vimentin, glutamate-aspartate transporter (GLAST), Postsynaptic Density Protein 95 (PSD-95) and neuronal filament (NF), and microtubule-associated protein 2 (MAP2), Aquaporin-4 (AQP4) and NeuN (see antibody information in **Supplemental Table S6**). These markers were distributed across five thermal channels and three fluorophore channels using the three different fluorescent dye and quencher pairs (Atto488-BHQ1, Atto565-BHQ2, and Atto647-BHQ3), with three targets assigned to each IR-activated imaging cycle.

Coronal mouse brain sections^38^ were cut on a cryostat, fixed, permeabilized, and stained with DNA-barcoded antibodies, followed by binding of the corresponding DNA thermal probes. Antibody staining concentrations are listed in Supplementary Table S2, and DNA-barcoded antibody probes were validated before multiplexed imaging (**Supplementary Figure S12-14**). After addition of AuBPs dispersed in imaging buffer, sequential IR irradiation pulses were applied to generate calibrated plasmonic heating spikes at 39, 48, 57, 65 and 72 °C (**Figure 6a**) and 5-plex protein imaging in the 647-nm channel was achieved (**Figure S15**). Each pulse activated one thermal channel, enabling repeated imaging of the same tissue region without iterative antibody staining or fluidic exchange.

**Figure 6.**
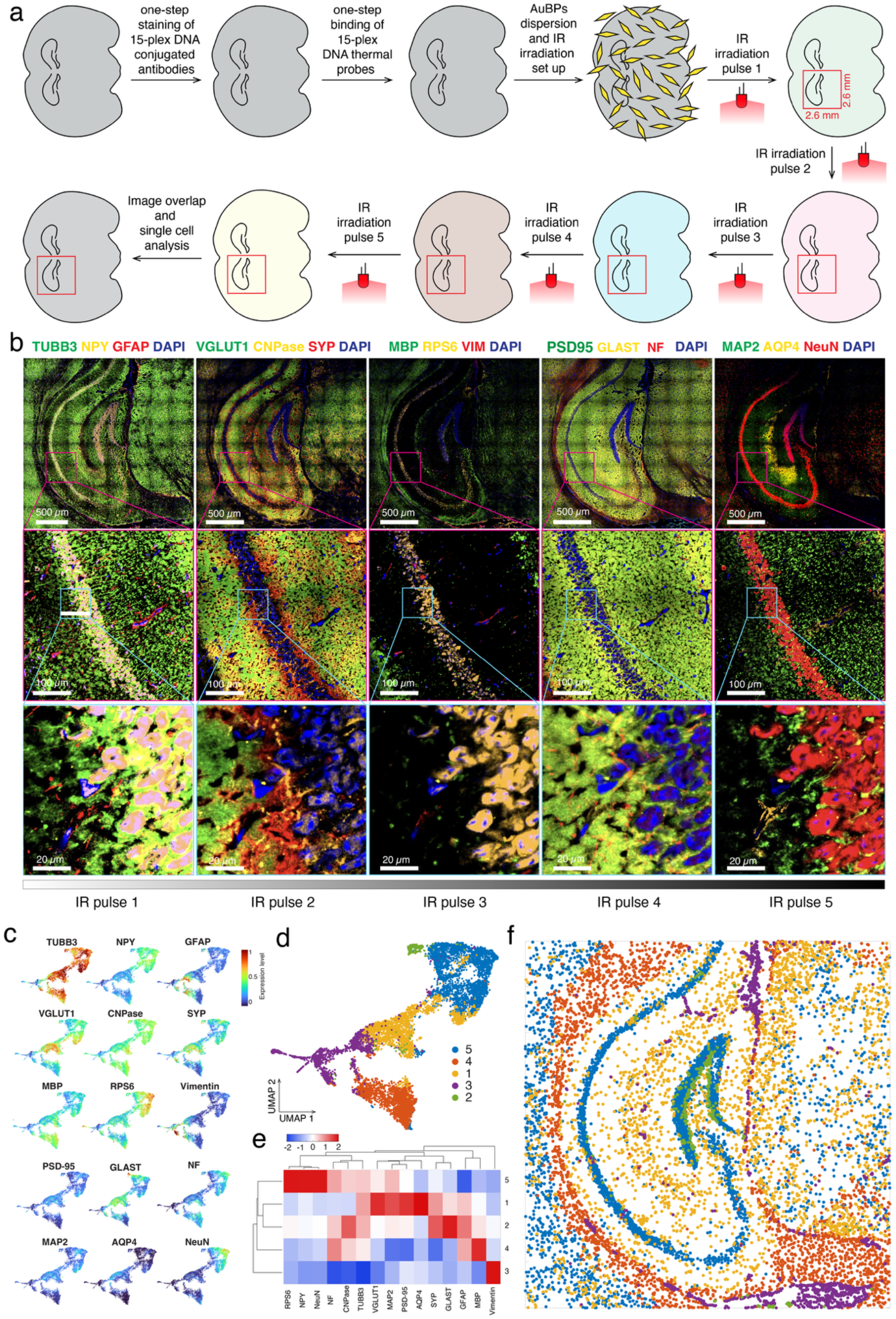
PHASER enables high-plex protein imaging in large mouse brain tissue sections and single cell multiparameter proteomic analysis. **(a)** Schematic workflow of PHASER for 15-plex protein imaging in mouse brain tissue slices. **(b)** Representative multiplexed fluorescence images of a mouse brain section acquired over five IR-activated imaging cycles corresponding to thermal channels of 39, 48, 57, 65, and 72 °C. Top row shows large-area hippocampal views, middle row shows enlarged regions indicated by orange boxes, and bottom row shows cellular-resolution zoomed-in regions indicated by magenta boxes. **(c)** UMAP visualization of single-cell protein-expression profiles from multiplexed mouse brain imaging, showing marker-specific expression distributions across the measured cell population. **(d)** Unsupervised clustering of single-cell protein profiles identifies five protein-defined cell groups. Their relative expression levels are shown as heatmap. **(e)** Heatmap showing marker enrichment z-score across the identified cell groups. **(f)** Spatial map of clustered cells projected back onto the tissue section, revealing the anatomical organization of protein-defined cell populations across the mouse brain region. 10641 total cells are analyzed from the 15-plex images.

We further used PHASER to resolve all 15 protein markers across an approximately 2.6 mm × 2.6 mm area to cover the hippocampus region of a mouse brain slice (**Figure 6b** and see individual protein images in **Supplemental Figure S16-18**). Large-area views captured anatomical organization across the hippocampal region, whereas 500 µm × 500 µm and 100 µm × 100 µm zoomed-in views showed cellular level marker distributions. The expected spatial patterns were readily observed: NeuN and MAP2 marked neuronal populations and dendritic structures, GFAP and GLAST highlighted astroglial features, aquaporin-4 labeled astrocyte-associated vascular and boundary structures, MBP and CNPase marked myelinated and oligodendroglial compartments, and VGLUT1, PSD-95 and NFL revealed synaptic and axonal organization. The preservation of these distinct staining patterns across sequential thermal channels indicates that PHASER can maintain tissue morphology and marker specificity during repeated plasmonic heating cycles. These results demonstrate that PHASER is scalable for high-plex, millimeter-scale spatial protein profiling in tissue samples.

### Single cell multiparametric spatial proteomic analysis enabled by PHASER

To determine whether PHASER-generated high-plex protein images could support spatial proteomic analysis at single-cell resolution, we segmented individual cells using DAPI nuclear signal with the quantified marker intensities as cytoplasmic signal in the registered 15-plex dataset with Cellpose-SAM^39, 40^ (see details in the Methods section) and obtained spatial expression profiles of 10641 cells (**supplemental Figure S19**). UMAP analysis of the resulting single-cell protein profiles revealed a structured expression landscape rather than a uniform distribution, demonstrating that the multiplexed PHASER measurements captured biologically meaningful molecular heterogeneity across the tissue (**Figure 6c**). Individual markers exhibited distinct but partially overlapping distributions in the UMAP space, consistent with the intermingled neuronal, glial, myelinated, and vascular-associated compartments of brain tissue.

Unsupervised clustering identified five groups with distinct protein expression signatures (**Figure 6d and e**). Group 5 was enriched in NPY, NeuN and RPS6, consistent with an NPY-positive neuronal population that may include hippocampal inhibitory interneurons^41^. Group 1 showed moderate enrichment of VGLUT1, AQP4 and TUBB3 without a single dominant marker, consistent with a mixed neuropil compartment containing both neuronal and astrocytic processes^42^. Group 2 was dominated by GLAST with a weaker contribution from GFAP, consistent with an astrocytic population defined primarily by glutamate-transporter expression^43, 44^. Enrichment of NF, MBP, and CNPase in cluster 4 was consistent with a myelin-rich compartment^45^. Group 3 was dominated by vimentin and may represent a structural, reactive-glial, or vascular-associated population. Overall, the distinct molecular signatures and spatial organization of clustered cells demonstrate that sequential photothermal activation, image registration, and single-cell quantification preserve both proteomic information and anatomical context. Together, the UMAP, clustering, heatmap, and spatial-projection analyses establish that PHASER-enabled high-plex imaging can generate quantitative, high-dimensional single-cell proteomic maps for studying the cellular and molecular organization of intact tissues.

## Discussion

PHASER provides a simple, rapid method for multiplexed fluorescence imaging in cells and tissues via light-controlled plasmic heating for the in situ activation of DNA thermal probe signal. The instrumentation only requires a low-cost LED light and gold bipyramid nanoparticles for convenient on-scope imaging. Melted imager and quencher DNAs remain at low (picomolar) concentrations, rendering their presence negligible in the imaging buffer and eliminating the need for fluidic exchange. Multiple rounds of signal switching can be achieved through consecutive irradiation of LED light-induced plasmonic heating, streamlining the workflow and reducing instrumentation complexity. We demonstrated the robustness of the methods for multiplexed imaging by resolving subcellular organelles and different neuron protein markers in brain tissues. The 5 rounds of signal exchange and imaging take less than 5 minutes to complete for a single field of view. This rapid workflow significantly enhances the throughput of cell and tissue sample imaging.

The multiplexity of PHASER can be significantly expanded through DNA probe design to improve the thermal channels and incorporate multiple fluorophore channels. Cooperativity^46^ can be implemented for DNA thermal probe design to narrow down the temperature range of DNA melting, significantly increasing thermal channels. By incorporating widely used 3∼5 fluorophore channels by different dye-quencher pairs^25^, the multiplexity can be improved by 3∼5 fold for 15∼25-plex imaging. With further enhanced multiplexity and extreme simplicity of PHASER in the future, complex brain circuits between neuron cells can be mapped to reveal brain functions. PHASER potentially can be applied to multiplexed super resolution imaging, such as DNA PAINT^12^. Traditional multiplexed super resolution with DNA-PAINT relies on the sequence specificity of different rounds of imagers, PHASER can be used to activate the blinking of DNA thermal probes in situ for rapid multiplexed super-resolution imaging after heating the desired temperature. PHASER can also be easily integrated with signal amplification strategy, such as SABER^13, 14^ and HCR^47^ to enhance the imaging signal and allow the detection of relatively low abundant targets.

Gold nanoparticle-based plasmonic heating with light is convenient to manipulate with a fluorescence microscope. The gold nanoparticle needs to be surface modified to be evenly distributed in the buffer, which is critical for even heating. Although this study focused on cultured cells and thin tissue sections, the approach has potential to be extended to thicker or three-dimensional samples. IR light is well suited for deeper optical penetration, and gold nanoparticles can potentially be distributed throughout gel-embedded or cleared tissue preparations.^48–50^ Overall, PHASER can provide a practical route toward rapid, accessible and scalable multiplexed protein imaging across diverse biological and clinical specimens.

## ACKNOWLEDGEMENT

This work is supported by the UF Start-up funding, UF McKnight Brain Institute Accelerator Award, NIH Maximize Investigators’ Research Award (R35GM165539) to F.H. The authors want to thank Dr. Boone Prentice for using their cryostats for brain tissue sectioning.

## AUTHORS CONTRIBUTIONS

E.X. designed and performed the study, analyzed the data. Y.C., Q.W., A.A.H., R. S.F. M. and M.Y.P. performed the experimental study. W.D.W., E.S., A.P.G., and Q.S. contributed to the supervision of the study. F.H. conceived, designed, and supervised the study and wrote the manuscript from all author’s input. All authors edited and approved the manuscript.

## DATA AND MATERIALS AVAILABILITY

The published article and the supplemental figures and tables include the datasets generated or analyzed during this study are available on Zenodo (DOI link: 10.5281/zenodo.21827326).

